# Targeting COL4A1-Related Small Vessel Disease: Repurposed Pharmacotherapies for Genetic Vasculopathies

**DOI:** 10.64898/2026.05.07.723462

**Authors:** Klaudia Kocsy, Harry Wilkinson, Zuzanna Sokolowska, Luke F Bolger, Vedanth Kumar, An Yan, Favour Felix-Ilemhenbhio, Alisdair McNeill, Sanjay Jain, Tom Van Agtmael, Mimoun Azzouz, Arshad Majid

## Abstract

COL4A1-related disorders cause early-onset stroke, intracerebral haemorrhage, visual impairment and kidney disease, often affecting children and young adults, yet no disease-modifying therapies exist. These disorders arise from pathogenic COL4A1 variants that disrupt type IV collagen and impair small-vessel integrity, leading to cerebral small-vessel disease and endothelial dysfunction.

We performed a mechanism-guided screen in human brain endothelial cells using a CRISPR-engineered COL4A1 p.G755R line and patient-specific COL4A1 p.G773R iPSC-derived endothelial cells. Simvastatin, L-carnosine, and XPD-101 restored impaired endothelial proliferation, migration, and other markers of endothelial function, including transendothelial electrical resistance (TEER). In a Col4a1^Svc/+^ mouse model, simvastatin increased pre-weaning survival, improved functional behaviour and reduced cerebral microhaemorrhage burden.

These findings identify mechanism-informed candidates that rescue COL4A1-mutant endothelial dysfunction in vitro, with simvastatin demonstrating in vivo efficacy, supporting prioritisation for further preclinical development.

## Introduction

The *COL4A1* gene encodes the α1 chain of type IV collagen, a crucial structural component of basement membranes that provides mechanical support and modulates cellular signalling across multiple tissues (1–3). Pathogenic variants in the *COL4A1* gene cause multisystem disorders affecting the cerebrovascular, muscular, ocular, and renal systems (2, 4, 5) ***(Figure 1A)***, collectively termed COL4A1-related disorders. Although the true prevalence of COL4A1-related disease is uncertain and likely underestimated, affected individuals experience substantial lifelong morbidity, and current care remains purely supportive (4, 6).

**Figure 1:**
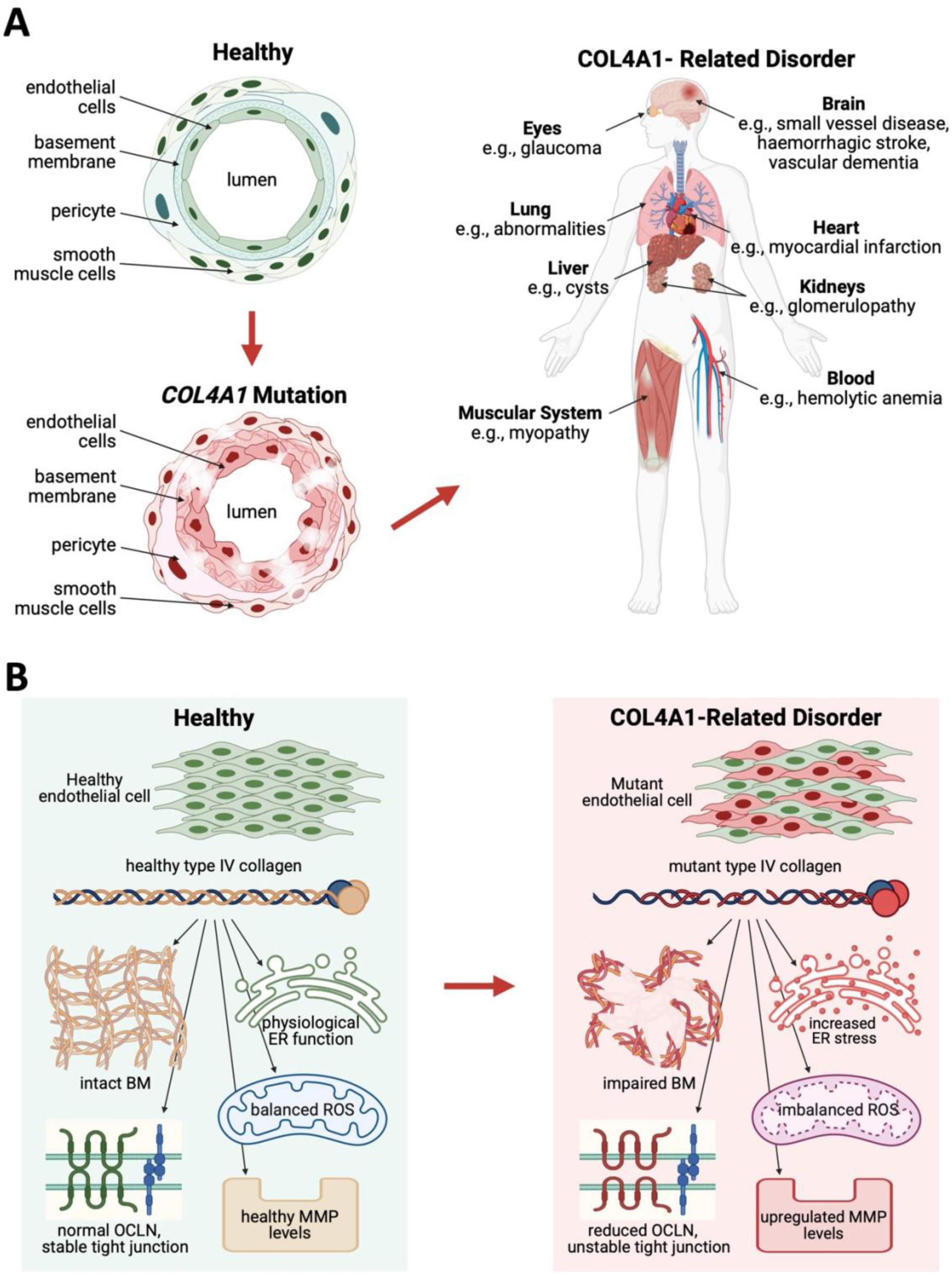
Wide-ranging clinical representation of COL4A1-related disorders. Healthy and diseased vessels illustrate basement membrane weakening, alongside a full-body view of the clinical spectrum (A). Structural differences between healthy and COL4A1-mutant collagen disrupt ER function, ROS levels, tight junction integrity, and MMP activity (B).

In the brain, *COL4A1* mutations lead to cerebral small vessel disease (CSVD) by disrupting the vascular basement membrane, weakening small vessels and predisposing to ischaemic and haemorrhagic strokes, as well as cognitive impairment (7–9). The collagenous abnormalities observed in CSVD closely mirror those in COL4A1- and COL4A2-associated disorders, underscoring the central role of basement membrane integrity in maintaining small vessel structure and function (9, 10). These tissue-level changes reflect more fundamental cellular dysfunctions that drive disease progression.

At the cellular level, COL4A1-related dysfunction is characterised by impaired cell proliferation and migration, as well as defective basement membrane assembly and integrity (1, 11) ***(Figure 1B)***. These structural defects are exacerbated by increased matrix metalloproteinase (MMP) activity, which further degrades the extracellular matrix, worsens tissue injury, and disrupts the blood-brain barrier (BBB) (12). Disorganisation of occludin and tight junctions compromises endothelial barrier function, thereby increasing vascular fragility (12). In parallel, COL4A1 mutations elevate reactive oxygen species (ROS) and induce chronic endoplasmic reticulum (ER) stress through intracellular accumulation of misfolded collagen, thereby contributing to progressive cellular dysfunction and multi-organ pathology (5, 12, 13).

Most pathogenic *COL4A1* variants are glycine substitutions within the triple-helical domain, which destabilise the collagen IV network and cause intracellular accumulation of misfolded protein, triggering chronic ER stress and oxidative injury in multiple organs (1, 4, 5, 13). These maladaptive stress responses, together with MMP-driven basement membrane degradation, are thought to be central drivers of endothelial dysfunction and small-vessel fragility in *COL4A1*/*COL4A2*-related small-vessel disease (4, 12, 13).

More than 850 cases of COL4A1-related disorders have been reported in the Gould Syndrome Foundation registry, while public variant databases catalogue approximately 3000 COL4A1 variants, many of which lack definitive clinical annotation. This genetic and phenotypic heterogeneity complicates variant interpretation and limits diagnostic classification, thereby likely underestimating the true burden of disease. These disorders are associated with severe, lifelong morbidity, yet management remains largely supportive, focused on symptom control and prevention of complications (4). The absence of disease-modifying therapies and the limited progress in therapeutic development highlight a critical need for strategies that directly target the underlying cellular mechanisms of disease.

To address this gap, we focused on key endothelial dysfunctions associated with COL4A1 pathology. We evaluated 13 mechanistically selected compounds for their ability to rescue pathogenic phenotypes in COL4A1-mutant endothelial cells, including MMP-driven matrix remodelling, oxidative and ER stress, and endothelial barrier dysfunction ***(*Table 1*)***. These compounds were assessed across in vitro and in vivo models of COL4A1-related disease.

**Table 1:** Summary of tested compounds and their mechanism of action.

| Compound | Mechanism of Action | Rationale | Clinical status | Ref |
| --- | --- | --- | --- | --- |
| **AZD1236** | Inhibit MMPs to prevent excessive ECM degradation | MMPs are upregulated in COL4A1 mutations, promoting tissue injury | discontinuation in COPD trials; preclinical research for other use | (12, 14) |
| **Marimastat** |  |  | discontinuation in cancer treatment; preclinical research for other use | (12, 15, 28) |
| **AZM198** | Suppresses myeloperoxidase (MPO) | Reduces endothelial dysfunction & oxidative stress | Preclinical research | (17) |
| **Aspirin** | Antiplatelet (COX-1, P2Y12) inhibitors | Prevent thromboembolic events | FDA/EMA-approved | (18) |
| **Clopidogrel** |  |  | FDA/EMA-approved | (19) |
| **Ramipril** | ACE inhibitor | Enhances endothelial function | FDA/EMA-approved | (20) |
| **Simvastatin** | Lower cholesterol, stabilize endothelium, reduce inflammation | Improve vascular health | FDA/EMA-approved | (21) |
| **Atorvastatin** |  |  | FDA/EMA-approved | (22) |
| **Taurine** | Modulates NMDA receptors | Limits excitotoxicity | supplement | (23) |
| **Vitamin C** | Antioxidant defence, vascular regulation | Support vascular health | supplement | (24) |
| **Vitamin D3** |  |  | supplement | (25) |
| **L-carnosine** | Antioxidants, reduce ischaemia-induced oxidative damage | Direct tissue protection | supplement; pre-clinical research | (26) |
| **XPD-101** |  |  | pre-clinical research | (26) |

AZD1236 (14) and Marimastat (15) inhibit matrix metalloproteinases (MMPs), which can be excessively activated due to COL4A1 mutations (16), thereby degrading extracellular matrix (ECM) components and regulating adhesion molecules. AZM198 (17) suppresses myeloperoxidase (MPO), reducing endothelial dysfunction and oxidative stress in vascular inflammation. Aspirin (18) and clopidogrel (19) inhibit platelet aggregation (via COX-1 and P2Y12, respectively), preventing thromboembolic events. Ramipril (20) (ACE inhibitor) enhances endothelial function. Simvastatin (21) and atorvastatin (22) lower cholesterol, improve endothelial stability and reduce inflammation. Taurine (23) modulates NMDA receptors to limit excitotoxicity. Vitamins C (24) and D3 (25) support antioxidant defences and vascular regulation. L-carnosine and XPD-101 (26) (antioxidants) alleviate ischaemia-induced oxidative damage. XPD-101 is a novel conjugate of D-carnosine and N-acetylcysteine amide (NACA), designed to overcome carnosinase-mediated degradation and enhance antioxidant and anti-excitotoxic capacity compared with L-carnosine (26, 27).

Here, we systematically evaluated 13 mechanistically selected compounds across COL4A1-mutant endothelial cell models and a Col4a1^Svc/+^ mouse model. Simvastatin, L-carnosine and XPD-101 emerged as lead candidates that rescued key endothelial phenotypes in vitro. Simvastatin, selected for in vivo evaluation based on its established clinical use and translational potential, additionally improved survival, behaviour and cerebral microhaemorrhage burden, supporting its prioritisation for further preclinical development in COL4A1-related small-vessel disease.

## Materials and Methods

### Cell Culture

Wild-type (WT) and CRISPR/Cas9-edited (COL4A1c.2263G>Ap.G755R) human cerebral microvascular endothelial cell (HBEC-5i) lines were kindly provided by Professor Tom Van Agtmael. These cells were cultured in Dulbecco’s Modified Eagle Medium: Nutrient Mixture F-12 supplemented with 40 μg/mL endothelial cell growth supplement and 10% foetal bovine serum.

Induced pluripotent stem cells (iPSCs) generated by Sanjay Jain (Washington University in St. Louis) from a COL4A1 patient (COL4A1c.2317G>Ap.G773R) and a healthy control (BJFF6) were differentiated into endothelial cells (iPSC-ECs), thereby providing patient-specific validation. iPSCs were maintained on Matrigel-coated vessels in mTeSR Plus medium and differentiated in two stages, first to mesodermal progenitors and then to endothelial cells, using StemCell Technologies differentiation kits according to the manufacturer’s instructions.

### Compound Treatment

HBEC-5i p.G755R and iPSC-EC p.G773R mutant cells were treated with the indicated compounds or vehicle control in their respective culture media for 72 hours under standard culture conditions (37°C, 5% CO₂) ***(*Table 2*)***. All compounds were prepared according to the manufacturers’ instructions, dissolved in the appropriate solvent, and sterilised using a 0.22 μm syringe filter before application. Following the treatment period, cells were washed with phosphate-buffered saline (PBS) and harvested for downstream analyses as described below. All drug treatments were performed in biological triplicate unless otherwise stated.

**Table 2:**
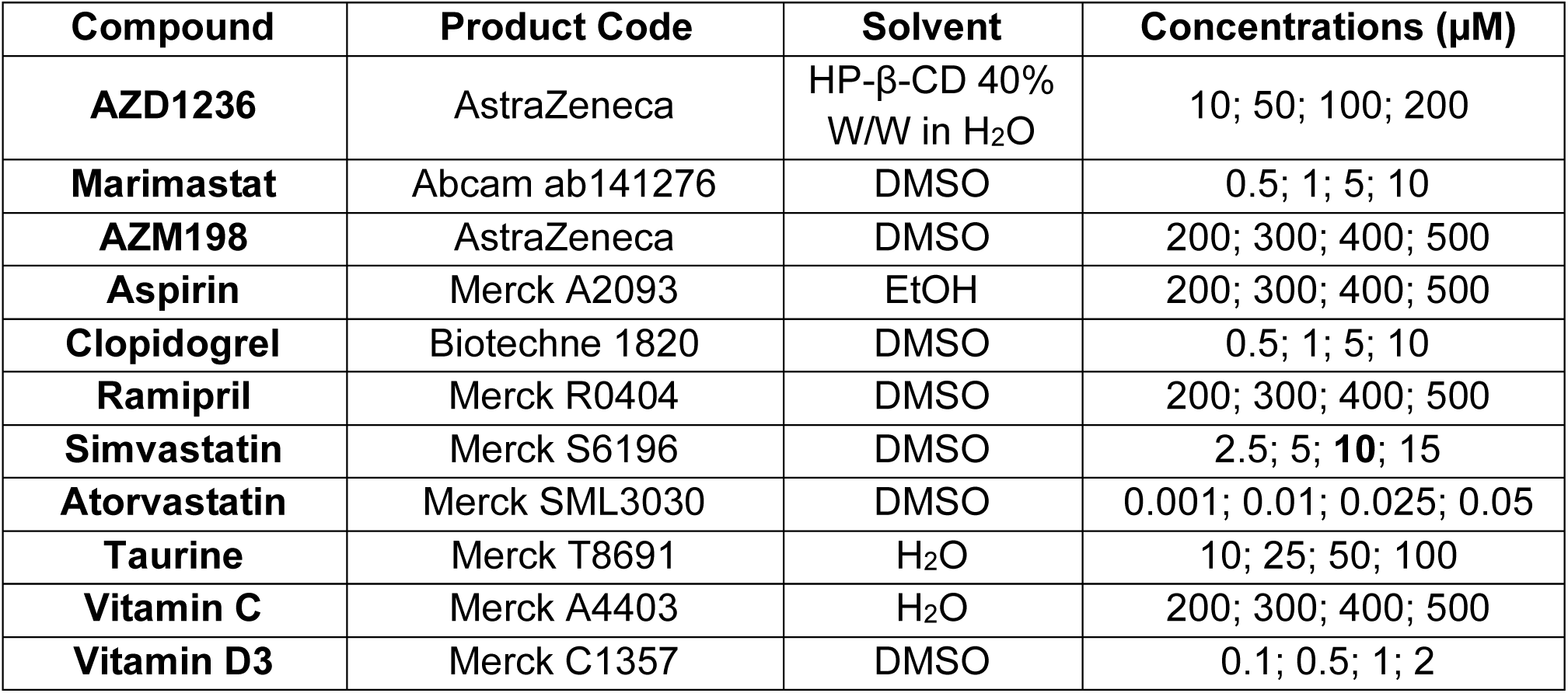
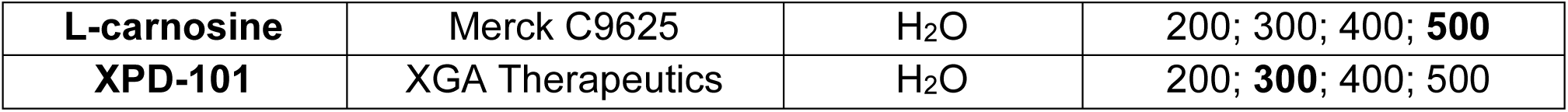
Summary of tested compounds, their solvents, and experimental concentrations. The highlighted concentrations were the best among the top three compounds.

### Cell Proliferation Assay

To assess proliferation, cells were seeded onto gelatine-coated 24-well plates at 1×10^4^ cells/well and treated continuously with the indicated compounds ***(*Table 2*)*** for a total of 72 hours before the first count and throughout the subsequent 5-day assay period. Cell numbers were then quantified every 24 hours for five days using a blinded haemocytometer count with trypan blue exclusion.

### Cell Migration Assay

The effect of compounds on cell migration was assessed using a scratch wound assay. Cell monolayers (>90% confluency) were scratched uniformly using the Incucyte Scratch Wound Maker (Sartorius). Migration was monitored for 24 hours, with images acquired every 2 hours using the Incucyte Live-Cell Analysis System (Sartorius), and wound closure was quantified using the associated Incucyte Cell Migration software.

Based on the proliferation and migration assays, the optimal concentration of each compound was defined as the dose that produced maximal phenotypic rescue without detectable cell death (also assessed with a cytotoxicity assay), and this dose was then used in all subsequent assays.

### Real-Time Quantitative PCR

Total RNA was extracted from compound-treated cells using the RNeasy Mini Kit (Qiagen) according to the manufacturer’s instructions. RT-qPCR was performed to assess expression of *COL4A1*, *OCLN*, *MMP2*, *MMP9*, ER-stress indicator genes (*ATF6*, *ERN1*, *HSPA5*, *XBP1*) and ROS-detoxifying genes (*SOD1*, *CAT*, *GPX1*). In HBEC-5i cells, target gene expression was normalised to the housekeeping gene *GAPDH*, whereas in iPSC-ECs, expression was normalised to *TMBIM6* (*29*). Data were analysed using the ΔΔCt method and expressed relative to wild-type cells.

The qPCRBIO SyGreen 1-Step Go Lo-ROX Master Mix (PCR Biosystems) was used, and data acquisition was performed using the CFX384 C1000 Touch Thermal Cycler (Bio-Rad).

### ELISA

To quantify protein levels of COL4A1, MMP2 and MMP9, cells were treated with compounds for 72 hours. For COL4A1 measurement, treated cells were lysed using immunoprecipitation (IP) lysis buffer supplemented with phosphatase and protease inhibitors, and cell lysates were analysed using a COL4A1-specific ELISA kit (Novus, NBP2-75870) according to the manufacturer’s instructions.

To assess secreted MMP2 and MMP9, after the initial 72-hour treatment, the culture medium was replaced with phenol red-free, serum-free medium and cells were incubated for a further 24 hours to minimise serum-derived MMPs and enrich for actively secreted forms. Conditioned media were then collected and analysed for total MMP2 (Thermo Fisher Scientific, KHC3082) and MMP9 (Thermo Fisher Scientific, BMS2016-2) following the manufacturer’s protocols.

For all ELISAs, samples were diluted 1:2 and measured in technical duplicates. Analyte concentrations were interpolated from standard curves according to the manufacturers’ instructions and then normalised to values obtained in wild-type cells. Absorbance was recorded using a PHERAstar plate reader.

### ROS production

Reactive Oxygen Species (ROS) were measured with the Cellular ROS Assay Kit (Abcam, ab186027) according to the manufacturer’s instructions (Ex/m = 520/605 nm).

### Collagenase Activity

Following compound treatment, cells were lysed in IP buffer, and total protein concentration was determined using the bicinchoninic acid (BCA) assay. Cell lysates were then subjected to a fluorometric collagenase activity assay to assess MMP activity using the EnzChek Collagenase Assay Kit (Thermo Fisher Scientific, E12055) according to the manufacturer’s instructions. Fluorescence was measured on a PHERAstar plate reader at 495 nm excitation and the corresponding emission wavelength, with readings acquired every 5 minutes over a 2-hour period.

### Cytotoxicity Assay

To evaluate compound-induced cytotoxicity, the viability of adherent cells was assessed using the Crystal Violet Assay Kit (Abcam, ab232855) according to the manufacturer’s instructions (570 nm).

### TEER (Transendothelial Electrical Resistance)

Transendothelial electrical resistance (TEER) was measured using an EVOM2 voltohmmeter (World Precision Instruments) with STX2 chopstick electrodes on gelatine-coated inserts (Thincert, 0.4µM pore diameter, Greiner Bio-One 662641). Compound-treated iPSC-ECs were seeded at 20,000 cells/insert and allowed to reach confluence. TEER was recorded at 48 h in culture medium after plates were equilibrated to room temperature for 15 minutes. Values represent the mean of three technical replicate measurements per insert, calculated by subtracting the mean blank insert resistance and multiplying by the insert surface area (0.336 cm²) to obtain Ω·cm², in line with MIRTA recommendations (30).

### iPSC maintenance, EC differentiation, and validation

Two iPSC lines (BJFF6 wild type and *COL4A1* c.2317G>A; p.Gly773Arg) were generated in collaboration with Professor Sanjay Jain (Kidney Translational Research Centre, Washington University). These were cultured on Matrigel (8.7 µg/cm²) in mTeSR Plus medium.

Endothelial cells were derived using the STEMdiff^TM^ Endothelial Differentiation system according to the manufacturer’s protocol, with sequential mesoderm induction (#05220), endothelial induction (#08005), and endothelial expansion (#08005). Differentiated endothelial cells were expanded up to passage 4 in expansion medium on 0.1% gelatine under standard culture conditions, and identity was confirmed by expression of endothelial markers PECAM1, VE-cadherin, and vWF.

### Mouse experiments

Mice were housed under specific pathogen-free conditions in individually ventilated cages on a 12 h light/12 h dark cycle, with ad libitum access to standard chow and water.

Col4a1^Svc/+^ mutant mice were used to assess the in vivo effects of simvastatin on survival and cerebrovascular outcomes. Simvastatin was administered via the drinking water at an estimated dose of 20 mg/kg/day. Pregnant dams received simvastatin from mating through gestation and lactation, and offspring continued treatment until three months of age, when nesting behaviour was assessed, and mice were sacrificed for endpoint analyses.

Nesting behaviour (n=7 per group) was assessed blindly at 3 months of age using a standard home-cage nesting assay, in which mice were provided with a pre-weighed nestlet and nest quality was scored the following day on a semi-quantitative scale (Deacon scale; 1-no nesting; 5-optimal nest).

Perls’ Prussian blue staining for intracerebral haemorrhage was performed on fixed brain sections using a standard iron detection protocol, and blinded microscopic evaluation of hemosiderin-positive foci (n=4 per group) presented as percentage of full brain slices.

All procedures complied with institutional and national animal welfare regulations and were approved by the relevant ethics committee.

### Statistics

All statistical analyses were performed using GraphPad Prism (version 10). For comparisons between two groups, the unpaired Student’s t-test was used, and for three or more groups, one-way analysis of variance (ANOVA) with Dunnett’s multiple comparisons test was applied. Data are presented as mean ± standard deviation (SD), with statistical significance set at 95% confidence (∗p ≤ 0.05, ∗∗p ≤ 0.01, ∗∗∗p ≤ 0.001, ∗∗∗∗p ≤ 0.0001). The reported “n” values refer to independent experiments (biological replicates).

Further details on experimental repeats and specific statistical tests are provided in the corresponding figure legends.

## Results

### Differences in wild-type (WT) and mutant (p.G755R) endothelial cells

We first confirmed that the COL4A1 p.G755R mutation causes marked endothelial dysfunction in HBEC-5i cells.

Compared with wild-type cells, mutant endothelial cells exhibited significantly reduced *COL4A1* mRNA expression (0.5-fold, p<0.0001; ***Figure 2A***). and lower protein levels (0.6-fold, p<0.0001; ***Figure 2B, C***), normalised to laminin, a stable protein in ECs associated with COL4A1-related disorders (5).

**Figure 2:**
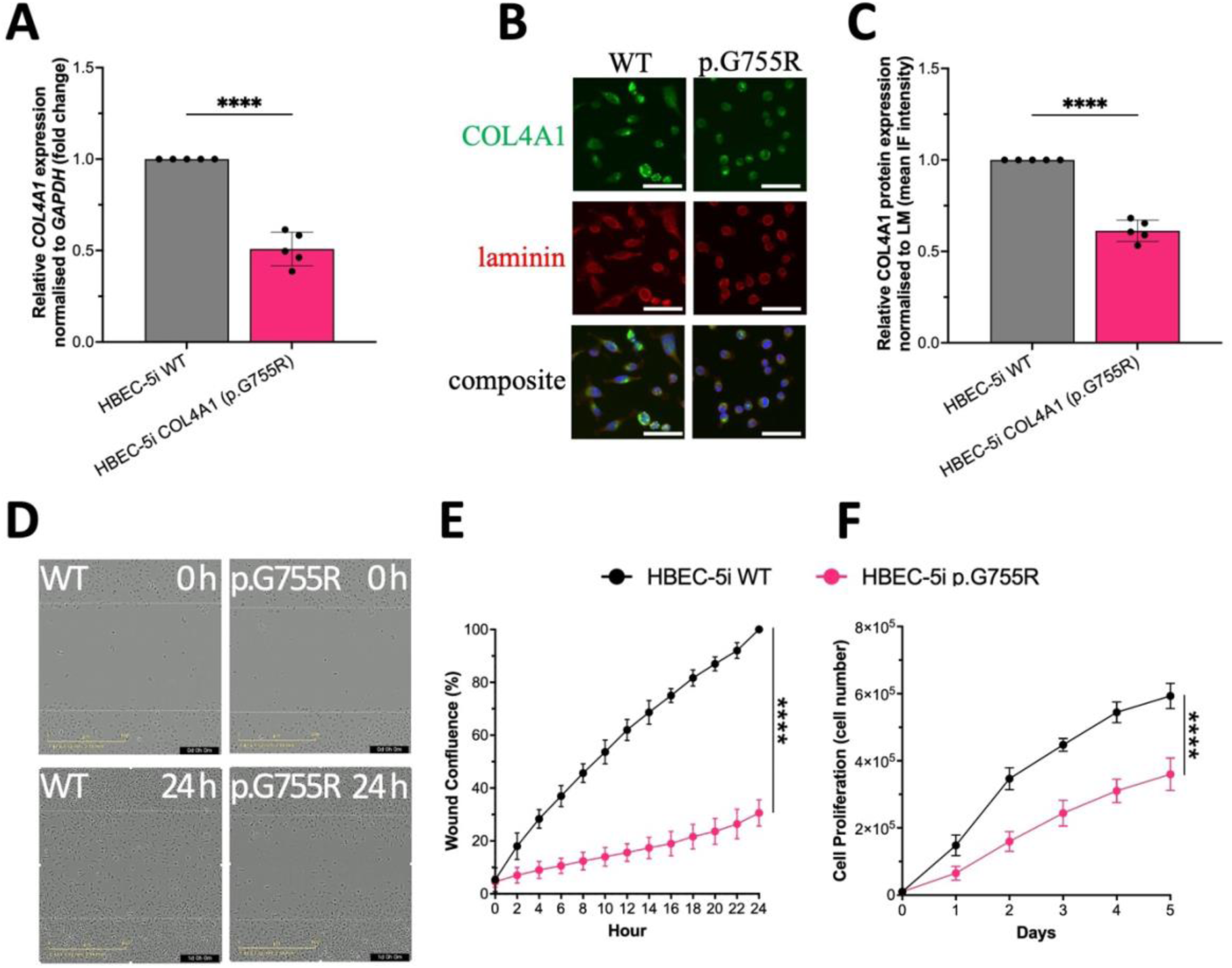
Mutant and wild-type endothelial cell characteristics. The expression of the COL4A1 gene and protein was markedly reduced at both the RNA (A, measured by Rt-qPCR) and protein levels (B-C) in the mutant cells, as measured by IF staining and normalised to laminin. Wound-healing and proliferation assays demonstrate reduced migration (69.4%, D-E) and proliferation (39.33%, F) in mutant HBEC-5i cells compared with WT. Results are presented as mean ±SD of 3 biological repeats (n=3) and analysed using a unpaired t-test; p-values: ***<0.0002; ****<0.0001.

Moreover, functional assays confirmed substantial phenotypic differences. In a wound-healing migration assay, mutant cells exhibited a 69.4% reduction in wound confluency compared to wild-type cells (p<0.0001; 30.6% of WT; ***Figure 2D, E***). Blinded cell count-based proliferation assays over five days revealed a 39.33% reduction in growth in mutant cells (p<0.0001; ***Figure 2F***).

These data demonstrate that the p.G755R mutation impairs key endothelial functions, establishing a quantitative baseline for evaluating pharmacological rescue in subsequent studies.

### Targeted drug screening in HBEC-5i cells

We next performed a mechanism-guided screen to prioritise compounds with the greatest potential to rescue these endothelial phenotypes.

We evaluated 13 repurposed or novel molecules at literature-supported concentrations, targeting pathways implicated in stroke and vascular integrity ***(*Table 1*)***.

Cell viability assays showed no detectable cytotoxicity at the tested doses, as measured by the Crystal Violet assay.

Results summarised in a heatmap format ***(Figure 3A)***, where each compound was ranked in each assay (1=phenotype closest to WT, 13=poorest), identified simvastatin, L-carnosine, and XPD-101 as the top three performers across assays.

**Figure 3:**
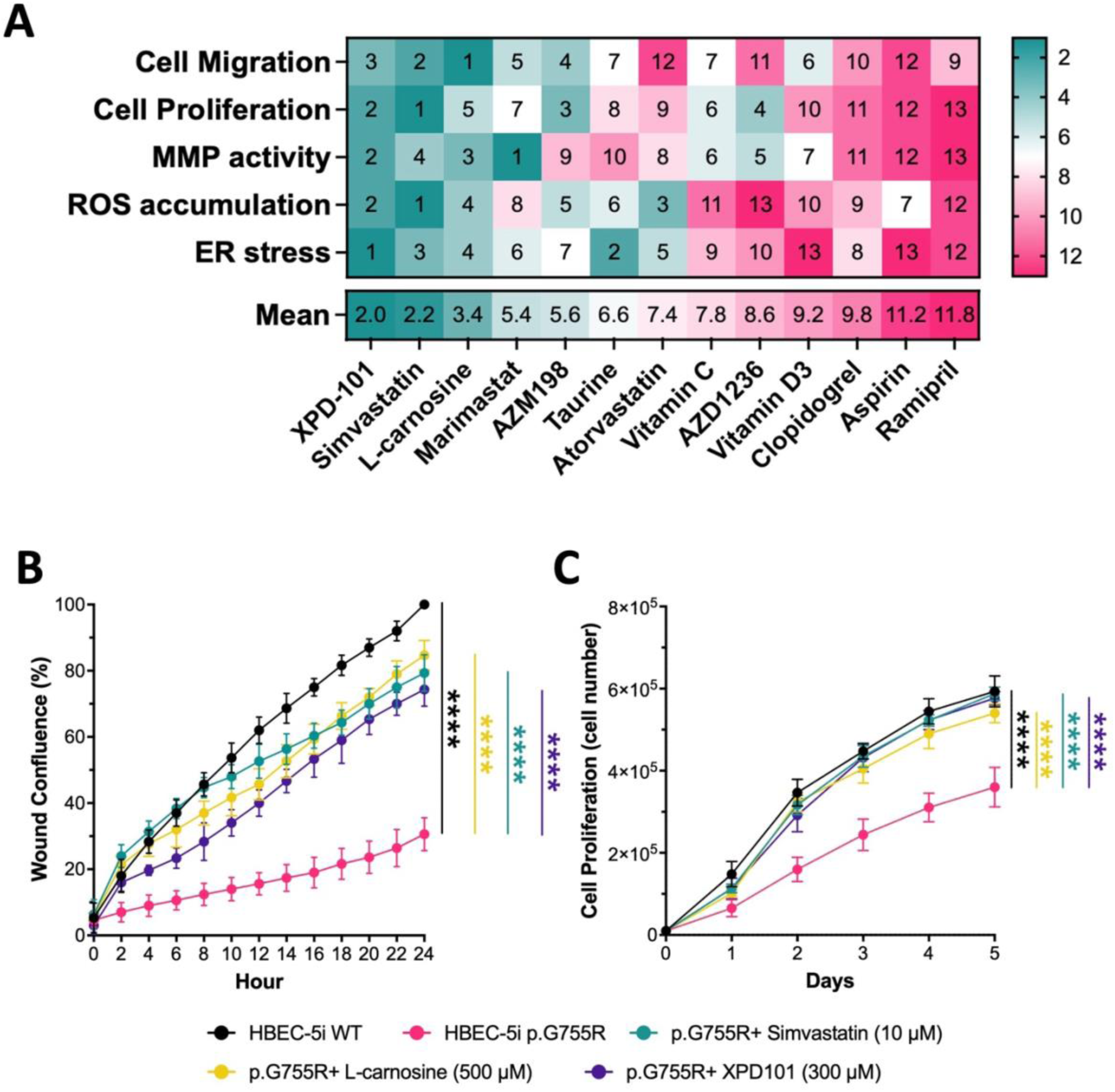
Top Compounds Identified by Phenotypic Screening Restore Cell Migration and Proliferation. Heatmap representation (A) of the efficiency of the 13 tested compounds. Each compound was assigned a rank in each assay, with 1 indicating that the mutant-treated cells’ phenotype closely resembled WT, and 13 indicating the poorest performance. The bottom row displays the mean rank for each compound across all assays, highlighting XPD-101, simvastatin, and L-carnosine as the top three performing compounds. Simvastatin, L-carnosine, and XPD-101 rescued cell migration (B), as measured by a scratch assay (Sartorius Incucyte). Similarly, these compounds restored cell proliferation (C), assessed every 24 hours for 5 days. Results are presented as mean ±SD of 3 biological repeats (n=3) and analysed using one-way ANOVA with Dunnett’s multiple comparison test; p-values: ***<0.0002; ****<0.0001.

### Restoration of cell migration and proliferation

To evaluate whether the lead compounds rescue core endothelial dysfunction, we assessed their effects on migration and proliferation in COL4A1 p.G755R mutant HBEC-5i cells.

After 72 hours of treatment with simvastatin (10 µM), L-carnosine (500 µM), and XPD-101 (300 µM) effectively restored p.G755R mutant cell migration ***(Figure 3B)*** and proliferation ***(Figure 3C)*** to near wild-type levels. In the scratch assay, migration increased to 79.3% of wild-type levels with simvastatin, 84.7% with L-carnosine, and 74.3% with XPD-101 (all p<0.0001). Proliferation was rescued to 97.5% of wild-type with simvastatin, 77.1% with L-carnosine, and 92.8% with XPD-101 (all p<0.0001). Together, these results demonstrate the strong potential of all three compounds to reverse the impaired cellular function observed in COL4A1-mutant endothelial cells.”

### Restoration of ER stress, ROS production and MMP activity

To determine whether these lead compounds also correct downstream stress pathways, we next assessed their effects on ER stress markers, ROS production, and MMP activity in COL4A1-mutant endothelial cells.

After 72 hours of treatment, simvastatin, L-carnosine, and XPD-101 significantly reduced the gene expression of ER stress markers *ATF6*, *ERN1*, *HSPA5*, and *XBP1*. These markers were elevated approximately 1.3-1.4-fold in mutant cells compared with wild type (all p≤0.0059) and were restored to or below wild-type levels (all p<0.0001; ***Figure 4A-D)***.

**Figure 4:**
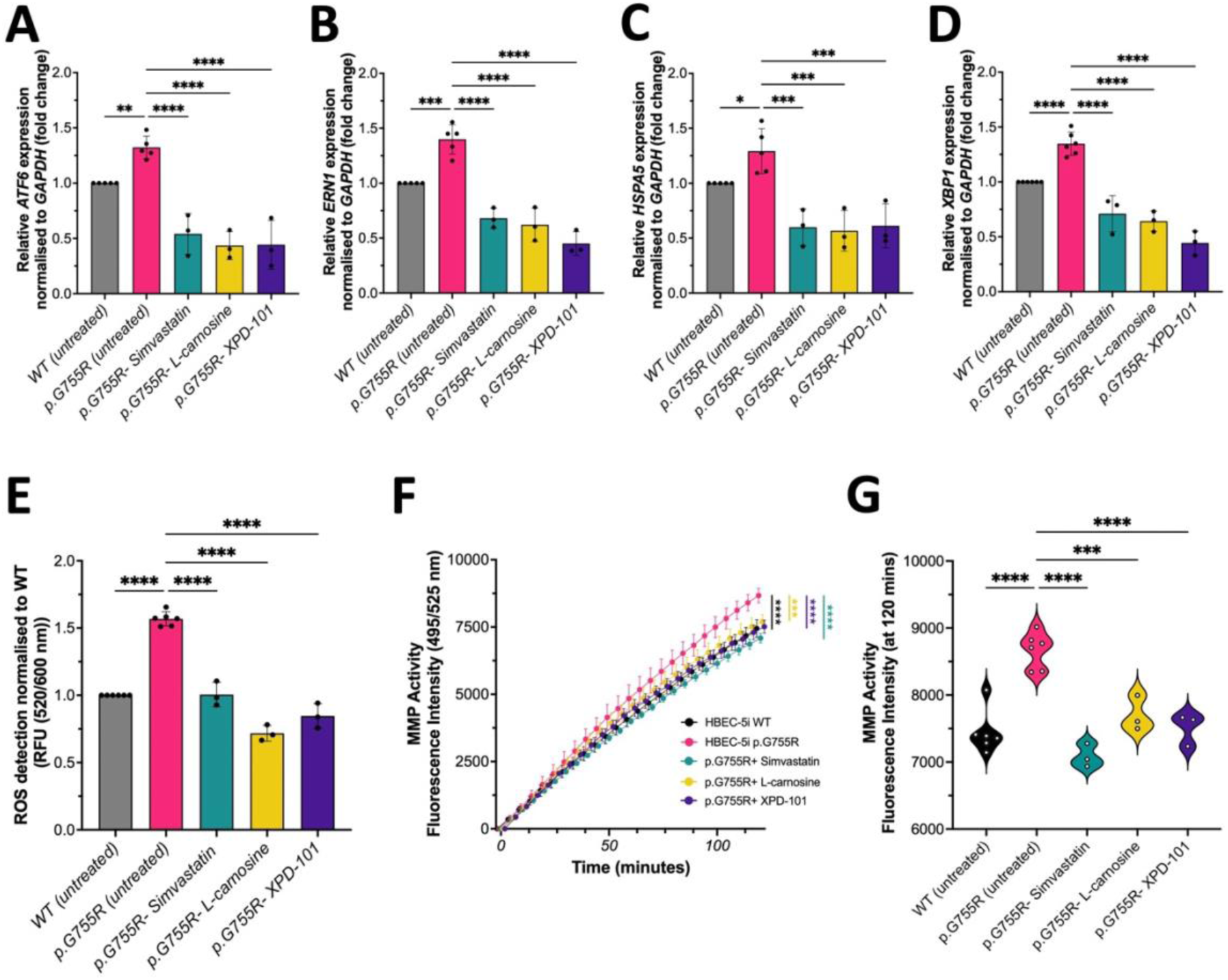
Treatment-Induced Changes in ER Stress, ROS Production, and MMP Activity. Decreased expression levels of the ATF6 (A), ERN1 (B), HSPA5 (C), and XBP1 (D) indicate a reduction in ER stress. Cellular ROS levels (E) and MMP activity (F) were restored, suggesting a rescue of the cellular phenotype. The violin plot (G) shows MMP activity at the 120-minute time point. Presented as mean ±SD of 3 or 5 biological replicates (n=3; 5) and analysed using one-way ANOVA with Dunnett’s multiple comparison test, p-value: *<0.0332; **<0.0021; ***<0.0002; ****<0.0001.

Meanwhile, the increased ROS signal in mutant cells (about 1.6-fold relative to wild type; p=0.0001) was normalised by all three compounds, with simvastatin restoring ROS to baseline levels and L-carnosine and XPD-101 reducing it to intermediate levels (all p=0.0001; fluorometric assay; ***Figure 4E)***.

Finally, the increased MMP activity in mutants (around 116% of WT) was reduced by each compound towards WT values, with all three treatments bringing MMP activity close to 100% (all p=0.0001, EnzChek Collagenase Assay; ***Figure 4F, G***).

Together, these findings show that the three small molecules effectively reduce ER stress, oxidative stress, and MMP imbalance in COL4A1 mutant endothelial cells.

### Effects on COL4A1, MMP2, and MMP9 Expression

To assess whether the lead compounds directly influence key matrix remodelling pathways in COL4A1-related disease, we next examined their effects on COL4A1, MMP2, and MMP9. Specifically, we measured mRNA levels by RT-qPCR ***(Figure 5 top row***) and quantified corresponding protein abundance by ELISA (***Figure 5 bottom row***).

**Figure 5:**
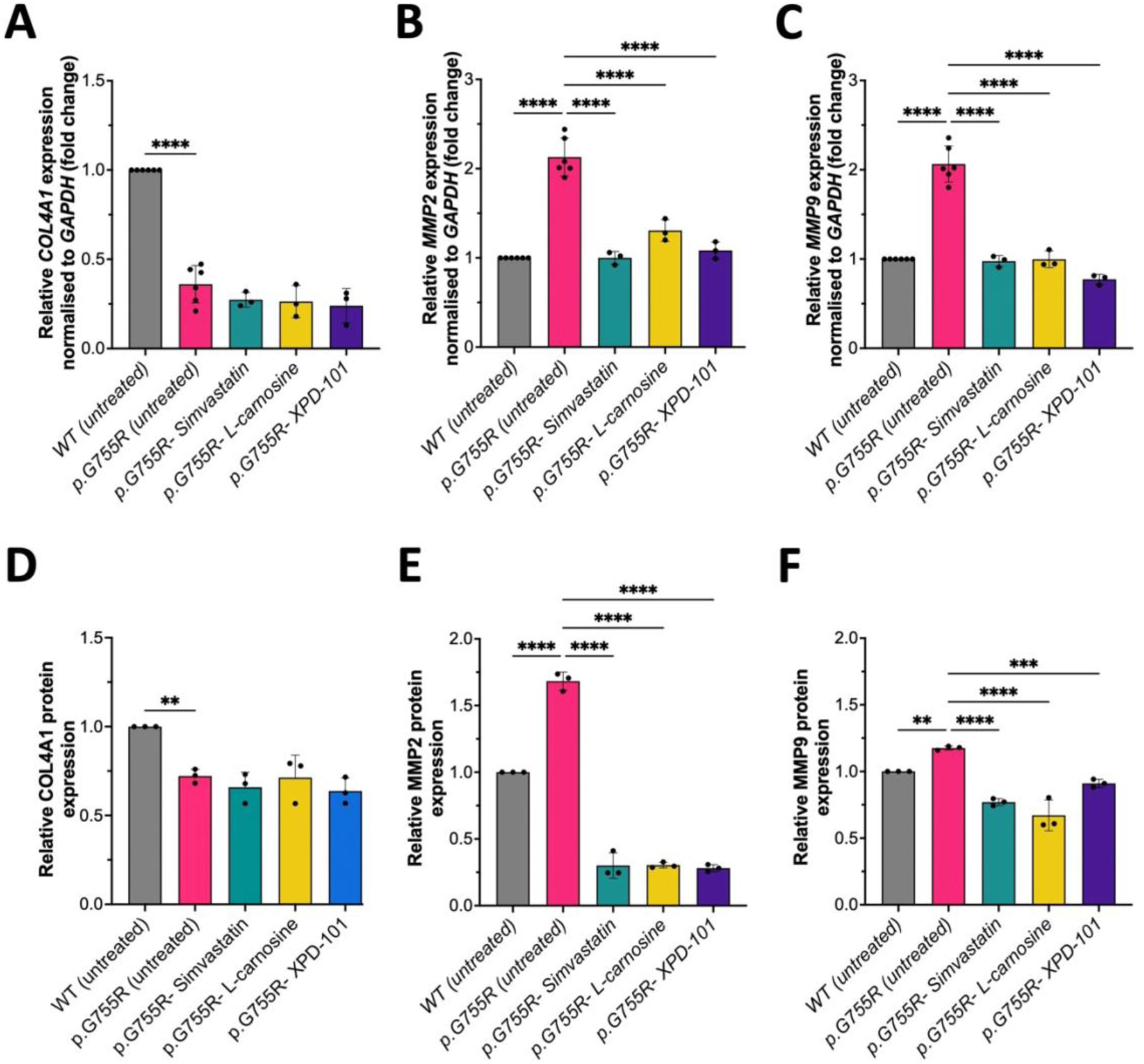
The mRNA (top) and protein (bottom) changes upon treatment. The COL4A1 expression levels (A, D) were not affected by any treatment, as measured by RT-qPCR (A) for mRNA and by ELISA (D) for protein levels. All three compounds reduced MMP2 (B, E) and MMP9 (C, F) levels measured by RT-qPCR and ELISA assays. Presented as mean ± SD of 3 or 5 biological replicates (n=3; 5) and were analysed using one-way ANOVA with Dunnett’s multiple comparison test, p-values: *<0.0332; **<0.0021; ***<0.0002; ****<0.0001.

*COL4A1* expression was markedly reduced in mutant endothelial cells at both the transcript (0.36-fold vs wild type, p<0.0001) and protein level (0.72-fold, p=0.0045), and none of the three compounds altered these levels (***Figure 5A, E***).

In contrast, *MMP2* and *MMP9* were strongly upregulated in mutant cells and were robustly normalised by treatment. *MMP2* mRNA increased to about 2.1-fold in mutants (P<0.0001) and was reduced to approximately 1.0-1.3-fold by simvastatin, L-carnosine, and XPD101 (all p<0.0001) (***Figure 5B)***. *MMP9* mRNA increased to around 2.1-fold (p<0.0001) and was brought back to near or below wild-type levels by all three drugs (p<0.0001, ***Figure 5C***). At the protein level, mutant cells showed elevated MMP2 (about 1.7-fold, p<0.0001) (***Figure 5E)*** and modestly increased MMP9 (1.18-fold, p=0.0097) (***Figure 5F)***, and treatment with each compound substantially lowered MMP2 and MMP9 proteins, with MMP2 reduced to roughly 30% and MMP9 to 67-91% of wild-type levels (all p≤0.005).

Together, these findings indicate that although the compounds do not alter COL4A1 abundance, they exert a strong inhibitory effect on MMP2 and MMP9 at both mRNA and protein levels, thereby supporting improved extracellular matrix stability.

### Validation in patient-specific iPSC-ECs

While CRISPR-engineered HBEC-5i cells provide a controlled platform to examine the effects of a single COL4A1 mutation, they do not fully reflect the broader genetic background and phenotypic variability observed in patients (31, 32). To provide patient-specific validation, we next tested the lead compounds in COL4A1 p.G773R patient-derived iPSC-ECs.

To determine whether the lead compounds also rescue functional deficits in patient-specific iPSC-ECs, we assessed cell migration and barrier integrity. Compared with wild-type cells, p.G773R mutant iPSC-ECs showed impaired migration, and all three lead compounds significantly improved this deficit (***Figure 6A)***. Migration increased to 87.3% of wild-type levels with simvastatin (p<0.0001), 64.7% with L-carnosine (p=0.0023), and 78.3% with XPD-101 (p<0.0001), demonstrating partial functional rescue of the migration phenotype in patient-derived endothelial cells.

**Figure 6:**
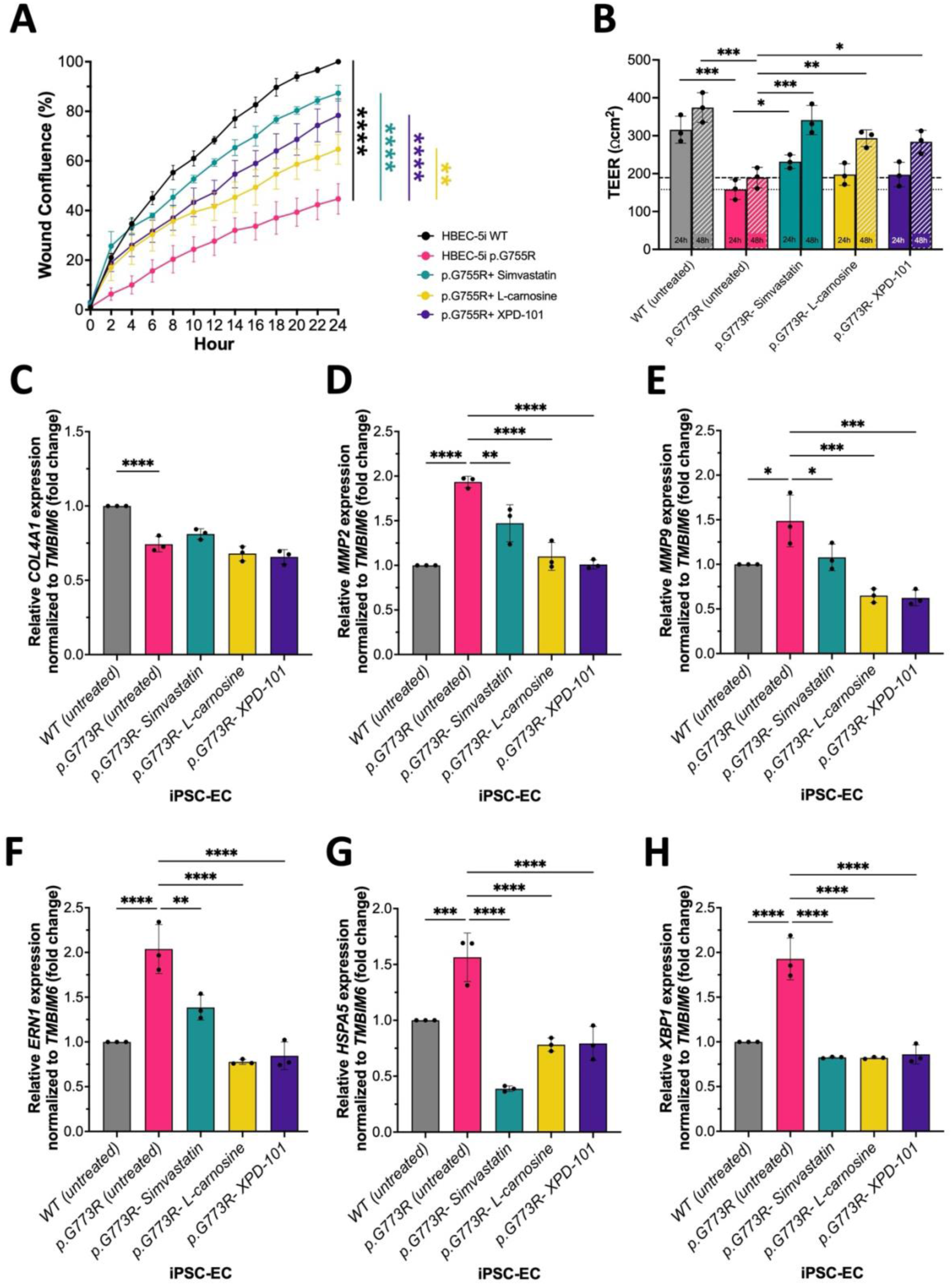
Beneficial effect of the top 3 compounds in patient-specific iPSC-ECs. Simvastatin, L-carnosine, and XPD-101 rescued cell migration in p.G773R mutant iPSC-ECs (A), as measured by a scratch wound assay (Sartorius Incucyte). The three small molecules increased TEER after 24 h and restored it close to wild-type levels at 48 h (B; dotted line: 24 h mutant baseline; dashed line: 48 h mutant baseline). These compounds did not modify the already reduced COL4A1 but normalised MMP2 and MMP9 expression and stabilised ERN1, HSPA5, and XBP1 levels towards wild-type values (C–H). Presented as mean ± SD of 3 biological replicates (n=3) and were analysed using one-way ANOVA with Dunnett’s multiple comparison test, p-values: *<0.0332; **<0.0021; ***<0.0002; ****<0.0001.

Because endothelial barrier integrity is essential for vascular stability, transendothelial electrical resistance (TEER) was measured as a functional readout of monolayer tightness and barrier function in iPSC-ECs. Mutant p.G773R cells showed significantly reduced TEER compared with wild-type cells after 24 hours in transwells (p=0.0002) and again at 48 hours (p=0.0001) (***Figure 6B)***. After 72 hours of compound pretreatment, TEER increased with the lead compounds after transfer to transwells. At 24 hours, simvastatin significantly increased TEER relative to untreated mutant cells (p=0.0341), restoring values to 73.4% of wild-type levels. At 48 hours, TEER was further improved by simvastatin (p=0.0006), reaching 91.4% of wild-type levels, while XPD-101 (p=0.00155) and L-carnosine (p=0.0088) also significantly increased TEER to 78.4% and 75.9% of wild-type levels, respectively.

To assess molecular changes, RT-qPCR was performed on key target genes. In mutant patient-specific cells, COL4A1 expression was reduced to 0.74-fold WT levels (p<0.0001), and treatment with the small molecules did not alter its expression (***Figure 6C)***.

*MMP2* was elevated to 1.93-fold in mutant cells (p<0.0001) and was partially normalised to 1.47-fold by simvastatin (p=0.0030), 1.10-fold by L-carnosine (p<0.0001), and 1.01-fold by XPD-101 (p<0.0001) (***Figure 6D)***. *MMP9* was increased to 1.49-fold in mutants (p=0.0107) and was reduced to 1.08-fold with simvastatin (p=0.0290), 0.65-fold with L-carnosine (p=0.0002), and 0.62-fold with XPD-101 (p=0.0002) (***Figure 6E)***.

ER stress-related markers were also elevated in mutant cells, with *ERN1* at 2.04-fold (p<0.0001), *HSPA5* at 1.56-fold (p=0.0007), and *XBP1* at 1.93-fold (p<0.0001) versus wild-type. Simvastatin reduced *ERN1* to 1.39-fold (p=0.0014), whereas L-carnosine and XPD-101 lowered it to 0.78-fold and 0.85-fold, respectively, compared to WT (both p<0.0001) (***Figure 6F)***. *HSPA5* expression decreased to 0.39-fold with simvastatin, 0.78-fold with L-carnosine, and 0.79-fold with XPD-101 (all p<0.0001) (***Figure 6G)***. *XBP1* was reduced to around 0.8-fold by all compounds (all p<0.0001) (***Figure 6H)***, restoring ER stress marker levels to near or below wild-type values.

These findings show that the three lead compounds not only normalise molecular stress markers in patient-specific iPSC-ECs, but also improve key endothelial functions linked to vascular maintenance, including migration and barrier integrity.

#### Simvastatin treatment in Col4a1^Svc/+^ mice

We next evaluated whether simvastatin confers survival and cerebrovascular benefits in Col4a1^Svc/+^ mice. Simvastatin was selected for in vivo evaluation based on its established clinical use and translational potential. It was delivered in the drinking water from mating until 3 months of age at 20 mg/kg/day (a safe dose in humans (33, 34).

Because Col4a1^Svc/+^ mutants are susceptible to early mortality (1, 13, 35), we evaluated survival at weaning age as a measure of viability, which increased from 75% to 90% of total births in vehicle- and simvastatin-treated litters, respectively (***Figure 7A)***. The ratio of heterozygous to wild-type offspring remained consistent between groups, indicating that simvastatin did not change the underlying genotype distribution.

**Figure 7:**
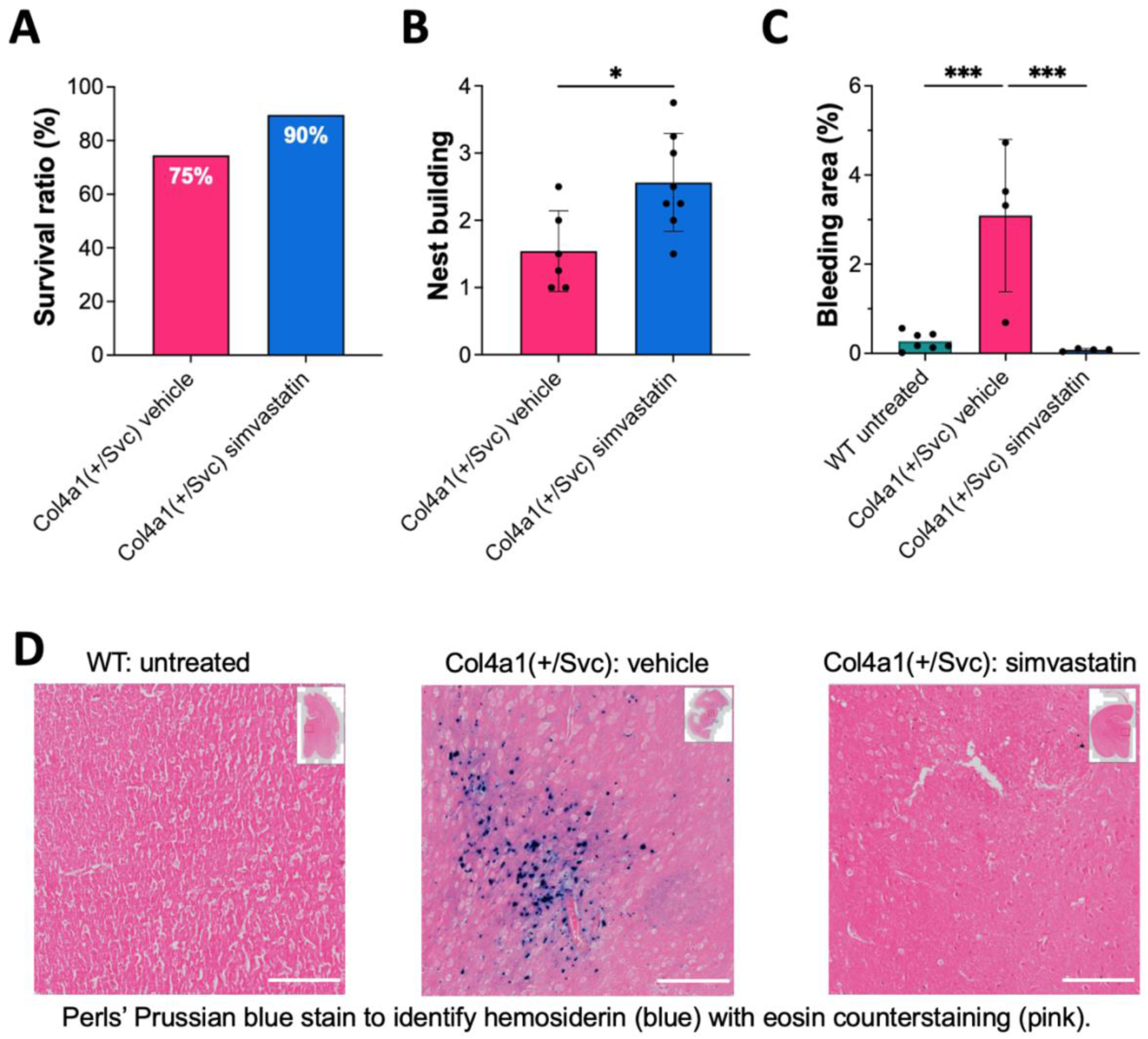
Simvastatin Improved Survival and Behavioural Outcomes and Reduced Microhaemorrhages in COL4A1^+/SVC^ Mice. Simvastatin treatment increased the proportion of surviving heterozygous pups compared to vehicle-treated littermates (A) and improved nest-building performance (B), as indicated by standardised nest scores (mean ± SD; unpaired t-test, p<0.032). Furthermore, simvastatin significantly reduced the cerebral bleeding area (expressed as % of the total brain section) (C) (mean ± SD; one-way ANOVA with Dunnett’s multiple comparison, p=0.0007). Representative brain sections stained with Perls’ Prussian blue (D) illustrate the reduction in haemorrhagic burden (Scalebar: 100 µM).

The pilot results also show a 66.3% increase in nest scores for simvastatin-treated mice compared with vehicle controls (SBE-β-CD; p=0.0164) (***Figure 7B)***. In this standardised assay, higher nest scores reflect more complex nest building, indicating improved functional behavioural performance.

Perls’ Prussian blue staining of brain sections from 3-month-old Col4a1^Svc/+^ mice revealed numerous hemosiderin deposits in vehicle-treated animals, consistent with previous descriptions of microhaemorrhage in this model (1, 35–37). Simvastatin-treated mice showed visibly fewer Perls’-positive foci. Quantitative analysis confirmed a significant reduction in bleeding area as a percentage of the total brain section. These results indicate that chronic simvastatin treatment reduces cerebral microbleeds in the Col4a1^Svc/+^ model (***Figure 7C, D)***.

## Discussion

In this study, we used a mechanism-guided screening strategy to identify compounds capable of rescuing core endothelial abnormalities associated with COL4A1-related disease. Across CRISPR-engineered HBEC-5i cells carrying the COL4A1 p.G755R mutation, patient-specific iPSC-derived endothelial cells carrying the p.G773R mutation, and an in vivo Col4a1^Svc/+^ mouse model, simvastatin, L-carnosine, and XPD-101 consistently improved key features of endothelial dysfunction. These included restored cell migration, proliferation, and transendothelial electrical resistance, together with reductions in elevated ER and oxidative stress and in MMP activity in vitro. In vivo, we focused on simvastatin, selected for testing because it combines robust in vitro efficacy with extensive clinical and safety data, and found that chronic treatment improved survival, enhanced nesting behaviour, and reduced the cerebral microhaemorrhagic burden in Col4a1^Svc/+^ mice.

Taken together, these findings support the concept that COL4A1-related small vessel disease is pharmacologically treatable by targeting convergent downstream mechanisms rather than directly restoring COL4A1 expression.

### Therapeutic rescue without restoring COL4A1 abundance

Across both our CRISPR-engineered p.G755R HBEC-5i line and the patient-derived p.G773R iPSC-ECs, we observe a convergent endothelial dysfunction phenotype characterised by reduced COL4A1 expression, impaired proliferation and migration, elevated MMP2/MMP9 activity, and increased ER and oxidative stress.

In these models, reduced proliferation, impaired migration, and lower transendothelial electrical resistance indicate a diminished capacity for endothelial repair and barrier function, key processes for maintaining vascular stability in development and adult tissues (12, 38–41). Our data, therefore, link glycine-substituting COL4A1 mutations to impaired vascular maintenance at the cellular level and are consistent with previous reports that collagen IV deficiency or COL4A1 mutations compromise endothelial and vascular cell proliferation and migration in vitro and ex vivo, rather than representing an artefact of our specific models (1, 5, 12, 13, 36).

Reduced COL4A1 levels in our models are particularly relevant, as defective collagen IV assembly and incorporation into the basement membrane are central features of COL4A1-related small-vessel disease and contribute to vessel fragility and haemorrhagic risk (4, 9, 42).

A key finding of this study is that pharmacological rescue occurred without recovery of COL4A1 expression itself. In both models, COL4A1 transcript and protein levels remained reduced after treatment, indicating that the beneficial effects of simvastatin, L-carnosine and XPD-101 are unlikely to reflect direct correction of mutant collagen biosynthesis. Instead, the observed rescue appears to arise from modulation of downstream pathological pathways, particularly endothelial dysfunction, oxidative and ER stress, and MMP-mediated matrix remodelling. This has important mechanistic and therapeutic implications. Most pathogenic COL4A1 variants disrupt triple-helix formation and activate chronic cellular stress responses (13, 43), making direct pharmacological restoration of normal COL4A1 biosynthesis difficult to achieve, especially in dominantly inherited structural disorders. Our findings, therefore, support targeting the consequences of the mutation rather than the mutation itself.

### MMP-driven matrix remodelling

Our data identify matrix metalloproteinase dysregulation as a particularly responsive and potentially central disease axis in COL4A1-mutant endothelial cells. Mutant cells showed increased total collagenase activity together with elevated MMP2 and MMP9 at both transcript and protein levels, closely matching a human COL4A1/A2 iPSC model in which excessive MMP2/MMP9 activity drives basement membrane loss and endothelial dysfunction (12). Given the established roles of MMP2/MMP9 in extracellular matrix degradation, vascular instability, and blood–brain barrier disruption, the magnitude and direction of these changes are consistent with a direct contribution to COL4A1-related vascular pathology (12, 44, 45).

These findings align with COL4A1–related pathology, in which basement membrane defects and vascular wall weakness are central, and with COL4A1/A2 iPSC models showing that MMP2/MMP9 upregulation drives loss of collagen IV and that pharmacological MMP inhibition can partially restore matrix integrity and endothelial function (12). Similar matrix remodelling processes have been implicated in arteriolosclerosis and white matter damage in sporadic cerebral small vessel disease, supporting the idea that MMP-mediated basement membrane degradation is a shared CSVD mechanism (7, 44, 45). Within this context, our observation that COL4A1 mutant endothelial cells display both enhanced MMP expression and increased collagenase activity reinforces MMP dysregulation as a core component of COL4A1-related small-vessel disease rather than a cell-line artefact.

Simvastatin, L-carnosine, and XPD-101 all normalised this molecular environment: each compound reduced MMP2/MMP9 expression and collagenase activity towards wild-type levels in both HBEC-5i and patient-derived p.G773R iPSC-ECs. These rescue effects are consistent with vascular studies showing that statins suppress MMP expression, reduce proteolytic activity and stabilise extracellular matrix in endothelial cells (46, 47). Clinically, simvastatin and related statins provide pleiotropic vascular benefits beyond lipid lowering, including modulation of endothelial-to-mesenchymal transition and inflammatory signalling, which may converge on reduced MMP activity and improved vascular stability (21). Together, these effects provide a plausible mechanistic link between MMP pathway suppression, improved endothelial function in vitro, and the reduced microhaemorrhagic burden observed with simvastatin in Col4a1^Svc/+^ mice.

Despite targeting matrix metalloproteinases, Marimastat and AZD1236 did not reproduce the functional rescue seen with our lead compounds. Both agents reduced MMP2 and MMP9 mRNA and protein levels, confirming on-target effects, but failed to improve migration, ROS, or ER stress markers, and had at most a marginal impact on proliferation. This disconnect between biochemical MMP suppression and lack of functional benefit indicates that direct MMP inhibition alone is insufficient to restore endothelial health in COL4A1-related small vessel disease, and that broader correction of interconnected stress and signalling pathways is required.

The comparison between simvastatin and atorvastatin further suggests that statin-mediated protection in COL4A1-mutant endothelium is drug-specific rather than class-wide. Both compounds similarly improved ER-stress markers and ROS, consistent with shared pleiotropic effects, but only simvastatin restored migration and more robustly enhanced proliferation and MMP normalisation. A key distinction is that simvastatin is more lipophilic, which may allow greater cellular and intracellular access and thus more effectively influence the cytoskeletal and adhesion mechanisms that underlie endothelial migration (48, 49).

Taken together, these observations indicate that MMP2/MMP9 dysregulation is a treatable component of COL4A1-related endothelial dysfunction, but that durable rescue requires agents such as simvastatin, L-carnosine and XPD-101, which couple MMP normalisation to relief of oxidative and ER stress and restoration of core endothelial functions. This integrated response may also be relevant to other inherited and sporadic forms of CSVD in which matrix remodelling and MMP imbalance play a central role.

### ER stress and oxidative stress as treatable targets

In addition to matrix remodelling, our study provides evidence that ER stress and oxidative stress are modifiable components of COL4A1 endothelial pathology. Mutant endothelial cells showed elevated expression of canonical ER stress markers, including *ATF6*, *ERN1*, *HSPA5*, and *XBP1*, as well as increased ROS production. All three lead compounds reduced these abnormalities substantially, with similar patterns observed in both HBEC-5i and patient-specific iPSC-EC models.

This is mechanistically consistent with COL4A1 biology: misfolded type IV collagen imposes a chronic proteostatic burden on the ER, driving sustained ER stress and UPR activation and secondarily increasing oxidative stress (1, 7, 12, 13, 43, 50, 51). Persistent activation of these pathways can impair endothelial homeostasis, exacerbate barrier instability, and promote abnormal matrix remodelling (13, 43).

Prior work in Col4a1/Col4a2 models has shown that targeting ER stress with chemical chaperones such as 4-phenylbutyrate (4-PBA) can reduce ER stress markers, improve collagen IV handling, and mitigate intracerebral haemorrhage or apoptosis, albeit with tissue- and allele-specific limitations (35, 52). Our findings extend this principle and show that clinically familiar or translationally actionable agents, including simvastatin and carnosine-based compounds, can also attenuate ER and oxidative stress, broadening the therapeutic landscape for COL4A1-related disease.

### Simvastatin may improve cerebrovascular outcomes

The Col4a1^Svc/+^ mouse is a well-established model of COL4A1-related small vessel disease, carrying a glycine-to-aspartate substitution that causes basement membrane defects, perinatal cerebral haemorrhages, reduced viability, and chronic cerebrovascular pathology (1, 5, 13).

In our study, chronic simvastatin treatment increased the survival of Col4a1^Svc/+^ offspring at weaning, improved nest-building scores and significantly reduced the proportion of microhaemorrhages. These findings suggest that targeting pathways modulated by simvastatin can translate into improved viability and functional behaviour, as well as reduced haemorrhagic burden in vivo.

They are broadly consistent with preclinical data showing that statins improve behavioural recovery and can reduce haemorrhage-related injury in experimental stroke and intracerebral haemorrhage models, largely through pleiotropic endothelial-stabilising, anti-inflammatory, and anti-oxidant effects (21, 22). Similarly, the protective effects of statins on haemorrhage-related injury have been observed in non-COL4A1 intracerebral haemorrhage models, where simvastatin and atorvastatin reduced haematoma volume, blood-brain barrier disruption, and tissue loss, while enhancing neurological recovery (53, 54).

Clinically, although very low LDL-cholesterol has been associated with increased ICH and haematoma expansion in observational cohorts (55, 56), randomised trials and meta-analyses indicate that standard-dose statin therapy does not consistently increase ICH risk and may improve survival and functional outcome after stroke (57–59). Our use of simvastatin at a human-relevant dose within the clinical range is aimed at harnessing these vascular benefits while avoiding excessive LDL-cholesterol lowering in a genetically haemorrhage-prone condition.

Taken together with our cellular data, this pilot supports simvastatin as a promising candidate for further preclinical studies in COL4A1-related small vessel disease, while also highlighting the need to define optimal dosing and timing to maximise benefit and minimise any potential haemorrhagic risk.

## Conclusion

Our data suggest that COL4A1-related endothelial dysfunction is pharmacologically tractable by targeting downstream pathogenic pathways rather than COL4A1 expression. Using a mechanism-guided screening approach, we identified simvastatin, L-carnosine, and XPD-101 as compounds capable of rescuing key cellular phenotypes, including impaired migration and proliferation, elevated ER and oxidative stress, and MMP dysregulation, across both engineered endothelial cells and patient-derived iPSC models. Together, these findings support the concept that downstream pathway modulation can enhance endothelial function and overall vascular health in COL4A1-related small-vessel disease. A strength of this work is the convergence of findings across CRISPR–edited HBEC–5i cells, patient–derived iPSC–ECs carrying a distinct COL4A1 variant, and an in vivo Col4a1^Svc/+^ mouse, supporting that the rescue reflects shared disease mechanisms rather than model–specific effects. Among the compounds tested, simvastatin showed the strongest translational potential, extending beyond in vitro rescue to improved survival, behavioural performance, and reduced microhaemorrhagic burden in Col4a1^Svc/+^ mice. Together with extensive clinical experience and safety data for statins (and, to a lesser extent, L-carnosine), these findings position simvastatin as a particularly attractive candidate for therapeutic repurposing in COL4A1-related small vessel disease, with XPD-101 providing a rationally designed tool compound to further refine pathway-targeted interventions.

Overall, this work provides a framework for mechanism-based therapeutic development in COL4A1-related disorders and supports further preclinical and translational studies to define optimal treatment strategies, dosing, and timing. More broadly, it suggests that pathway-targeted vascular stabilisation strategies developed in COL4A1-related CSVD may be relevant to other inherited and sporadic forms of small vessel disease characterised by basement membrane dysfunction and endothelial instability.

### Limitations

Our study has limitations. First, our in vitro work focused on endothelial monocultures with both glycine-substituting COL4A1 variants (p.G755R and p.G773R). We therefore did not directly assess the contributions of pericytes, vascular smooth muscle cells, or astrocytes, and the multicellular aspects of COL4A1-related pathology and treatment response may have been underestimated.

Second, the compounds were selected based on prior mechanistic rationale rather than identified through an unbiased high-throughput screen. While this targeted approach enabled pathway-focused interrogation, it may have excluded other potentially effective compounds.

Third, the in vivo component should be considered exploratory. We used a single Col4a1^Svc/+^ mouse line, a single simvastatin dose, and a single treatment window, with limited haemorrhage imaging endpoints. We also did not assess long-term safety or evaluate treatment effects in other affected organs, such as the eye, liver or kidney. Accordingly, these findings require confirmation in larger, powered studies.

### Future Directions

Future work should extend these findings into more complex multicellular systems. Multicellular blood-brain barrier models and brain organoids incorporating endothelial cells, pericytes, astrocytes, and neurons would allow direct assessment of barrier integrity, neurovascular coupling, and cell-cell interactions in response to COL4A1 mutations and candidate therapies. In parallel, in vivo studies should systematically explore dose-response relationships and treatment timing, and evaluate whether combination regimens, such as statin plus XPD-101, provide additive or synergistic benefit compared with monotherapy, ideally across multiple Col4a1 and Col4a2 mouse models that capture the broader mutational and phenotypic spectrum of COL4A1/2-related small vessel disease.

## Author Contributions

Experimental Design: K.K.; A.M.; T.V.A.

Experimental Work: K.K.; H.W.; Z.S.; L.F.B.; V.K.; A.Y.

Original Draft Preparation: K.K.

Review & Editing: K.K.; H.W.; F.F.-I.; T.V.A.; A.M.; S.J.; M.A.; A.M.

Visualisation: K.K.; A.M.

Resources/Ethical Approval: T.V.A.; A.M.; S.J.; M.A.; A.M.

Funding Acquisition & Project Supervision: A.M.

All authors have read and agreed to the published version of the manuscript.

## Declaration of generative AI and AI-assisted technologies in the writing process

While preparing this work, the author(s) used Perplexity AI and Grammarly AI writing Assistance to correct grammar and improve readability. After using these tools, the author(s) reviewed and edited the content as needed and take(s) full responsibility for the content of the publication.

## Research Publications and Copyright Policy

For the purpose of open access, the author has applied a Creative Commons Attribution (CC BY) licence to any Author Accepted Manuscript version arising from this submission.

## Figures and Graphs

Figures were created in BioRender. Graphs were generated in GraphPad Prism version 10.

